# Preservation of circadian rhythms by the protein folding chaperone, BiP

**DOI:** 10.1101/348078

**Authors:** Adam Pickard, Joan Chang, Nissrin Alachkar, Ben Calverley, Richa Garva, Peter Arvan, Qing-Jun Meng, Karl E. Kadler

**Affiliations:** Wellcome Centre for Cell-Matrix Research, Faculty of Science and Engineering, University of Manchester, Manchester, M13 9PT UK; Faculty of Biology, Medicine and Health, Manchester Academic Health Science Centre, and School of Mathematics, Faculty of Science and Engineering, University of Manchester, Manchester, M13 9PT UK; Division of Metabolism, Endocrinology, & Diabetes, University of Michigan, Ann Arbor, MI 48105 USA

**Keywords:** 4PBA, BiP, Collagen, Circadian rhythm, ER stress, PER2::Luc, SERCA, TUDCA, UDCA

## Abstract

ER stress and dysregulation of collagen synthesis are associated with progression of disease in cancer and fibrosis. Collagen synthesis is co-ordinated with the circadian clock, which curiously in cancer cells, is deregulated by ER stress. We hypothesised that interplay exists between circadian rhythm, collagen synthesis and ER stress in normal cells. Here we show that fibroblasts with ER stress do not demonstrate circadian rhythms in gene expression upon clock-synchronizing time cues. Conversely, overexpression of BiP or treatment with chemical chaperones strengthens the oscillation amplitude of circadian rhythms. The significance of these findings was explored in tendon, where we showed that BiP expression is ramped preemptively prior to a surge in collagen synthesis at night, thereby preventing protein misfolding and ER stress. In turn, we propose, this forestalls activation of the unfolded protein response in order for circadian rhythms to be maintained. Thus, targeting ER stress could be used to modulate circadian rhythm and restore collagen homeostasis in disease.

## Introduction

Circadian clocks are cell-autonomous time keeping mechanisms that occur in most tissues to optimize cellular activities in anticipation of varying demands on the cell during 24 hours ^1, 2^. One such function is control of translation and protein synthesis ^3^. In a recent study, Yeung and Garva *et al.* showed that the circadian clock regulates Sec61 translocon-dependent ER protein synthesis and membrane trafficking at each node in the secretory pathway of tendon fibroblasts *in vivo*^4^. The consequence is a daily surge of collagen-I production, which occurs at night in mice. Collagen-I is the major secreted protein of fibroblasts and, in tendon, accounts for 70% of the mass of the tissue where it occurs as extracellular fibrils^5^ that can extend the length of the tissue^6^. Collagens are large trimeric molecules that undergo extensive post-translational modification^7, 8^ and folding in the endoplasmic reticulum (ER, their site of synthesis) in order to generate a thermally-stable triple helix capable of assembling into fibrils. The extensive nature of posttranslational modification of collagen, coupled with a surge in synthesis at night, suggested that the cell might have mechanisms to protect against the accumulation of unfolded collagen (e.g. with the unfolded protein response, UPR) and subsequent ER stress, at times of high collagen synthesis. Excessive collagen synthesis occurs in fibrosis and solid tumours where the rate of synthesis exceeds turnover. The result is accumulation of a dense, stiff extracellular matrix that can be a positive fibroproliferative feedback signal to accelerate cell division and the synthesis of additional matrix^9^.

Type 1 collagen molecules (collagen-I) are composed of two **α**1 chains and one **α**2 chain encoded by the genes COL1A1 and COL1A2, respectively. These chains come together to form a triple helix, which is assisted by chaperone proteins. Both generic protein chaperones such as BiP/Grp78 and Grp94, and the collagen specific chaperone, Hsp47, aid the formation and stabilisation of the triple helix^10^. Chaperones also have signalling roles associated with their ability to bind newly synthesised proteins. In particular, BiP is known to govern the activation of the UPR^11^. When unfolded proteins accumulate in the ER, BiP is released from three key binding partners on the ER membrane; PERK, IRE1 and ATF6 each activate mechanistically distinct pathways. Combined, the response acts to reduce ER burden, through mRNA decay, and reducing translation, but also drive expression of protein chaperones. However, if the stress is not resolved these same pathways can also promote apoptosis^12^. It has been established that the response to ER stress is regulated in a circadian manner in cancer cells^13^ and that activation of UPR in cancer cells induces a 10 hour shift in circadian oscillation^14^; however it remains unknown how protein flux through the ER, and in turn how ER stress, regulates circadian rhythm in non-tumorigenic models. Given that collagen secretion is ramped from low levels during the day to high levels in the night in mice^4^, we hypothesised that the expression of these chaperones may be governed by circadian rhythm. In the experiments described below, we explored the connection between ER stress and circadian rhythm in fibroblasts actively secreting collagen.

## Results

### BiP protein levels are circadian clock rhythmic

BiP is known to interact with collagen in the secretory pathway^15^ and is proposed to aid the folding of collagen. As a first experiment, we showed that levels of BiP protein are rhythmic with a 24-hour period and its peak of expression precedes the peak of expression of procollagen-I (the precursor of collagen) protein (Fig. 1A). The circadian nature of BiP protein expression was also observed in mouse embryo fibroblasts (MEFs), isolated tail tendon fibroblasts, and tail tendon tissue (Fig. 1B, Supplementary Fig. 1). This result was the first indication that the circadian clock had a role in pre-emptively preventing ER stress by upregulating BiP in anticipation of a surge in collagen synthesis.

**Figure 1:**
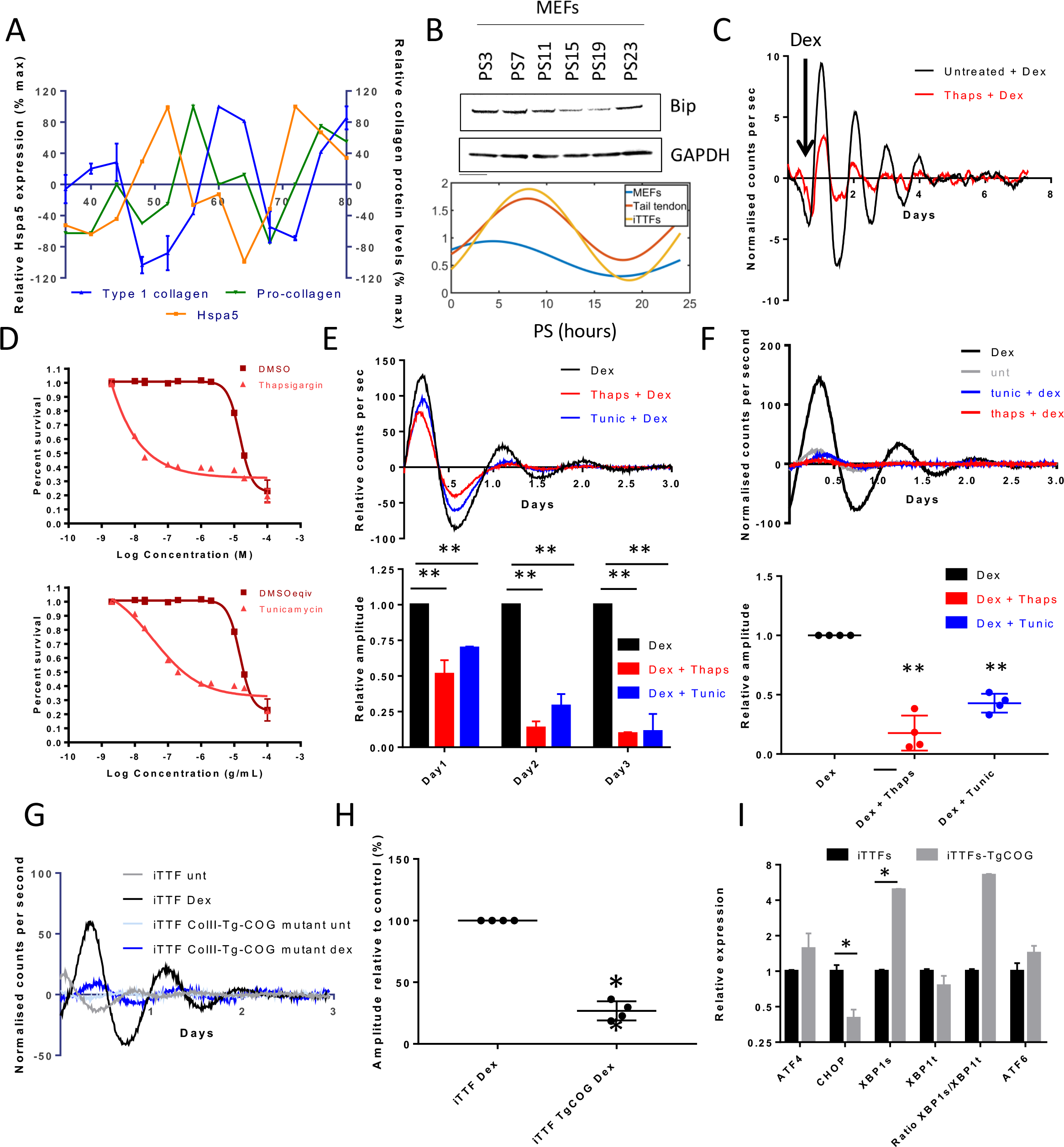
ER stress suppresses circadian rhythm. **A**) Hspa5 (BiP/GRP78) protein levels fluctuate over the course of 2 days in mouse tail tendons, BiP levels rise prior to the increased synthesis of collagen, indicated by pro-collagen specific peptides. **B**) BiP levels as assessed by western blot in MEFs over the course of 24 hours following synchronisation with 50% horse serum. Lower traces show the fluctuations in BiP protein levels in Tail tendons, iTTFs and MEFs as assessed by circwave ^43^ (see supplemental Fig. 1). **C**) The effects of ER stress induction on circadian rhythm in tail tendons from Per2::luc mice. **D**) Cell viability after 72 hours in response to the indicated doses of thapsigargin (Upper) and tunicamycin (lower). **E**) Representative luminescence traces (upper) following 1 hour treatment of 10 nM thapsigargin or 100 ng/mL tunicamycin which suppresses the induction of Per2::luc in response to dexamethasone, relative amplitude over 3 days are shown, n=3 (lower) **F**) Thapsigargin and tunicamycin are removed from Per2::luc cells after 5 hours treatment then synchronised with dexamethasone. Relative amplitudes in the first 24 hours are shown (lower chart, n=4). **G**) iTTF expressing mutated Thyroglobulin (ColII-Tg-COG mutant) have suppressed inherent and Dexamethasone-induced rhythms. Amplitudes of Per2::luc traces are quantified in **H**). **I**) Levels of spliced XBP1 (Xbp1s) indicate that the IRE1-XBP1 arm of the unfolded protein response has been activated in ColII-Tg-COG cells. **p<0.01. paired t-test.

### The presence of misfolded protein ablates circadian rhythm

To learn more about the effects of ER stress on the circadian rhythm, we treated ex vivo PER2::Luc clock reporter mice ^2^ tail tendons with thapsigargin to induce protein misfolding and ER stress^16^. When cells were subjected to dexamethasone to synchronise the circadian clock, the amplitude of rhythm was markedly reduced after thapsigargin treatment (Fig. 1C). Similarly, when immortalised tail tendon fibroblasts (iTTF) from Per2::luc mice were pre-treated for 1 hour with tunicamycin or thapsigargin, at doses which do not impact on cell growth (Fig. 1D), the amplitude of Per2 rhythm was suppressed (Fig. 1E).

This connection between ER stress and circadian rhythm was further exemplified in cells that were in the process of resolving ER stress prior to dexamethasone treatment. We used a 5-hour treatment with thapsigargin or tunicamycin to accumulate misfolded proteins in the ER and induce all arms of the UPR (Supplementary Fig. 2) prior to synchronisation of the circadian rhythm. This treatment regime eliminated the synchronisation effect of dexamethasone and forskolin (Fig. 1F, Supplementary Fig. 3), which suggested that cells undergoing ER stress do not have a functioning circadian clock. Similarly, in cultures with a pre-established rhythm treatment with thapsigargin flattened Per2:luc fluctuations (Supplementary Fig. 3).

To establish if misfolded proteins instigate suppression of circadian rhythms, we used a mutated thyroglobulin which misfolds in the ER, thereby causing ER stress^17^. When stably expressed in Per2::luc cells there was dramatic suppression of circadian rhythm (Fig. 1G and H) and induction of ER stress, particularly increasing the splicing of XBP1 (Fig. 1I). Thus, the fact that circadian rhythm is lost in fibroblasts expressing misfolded thyroglobulin shows that the loss of rhythm is the direct consequence of accumulated misfolded protein.

### Suppression of the secretory pathway blocks circadian rhythm by induction of ER stress

The effect of ER stress induction on collagen secretion was examined by western blot analysis of the conditioned medium of thapsigargin or tunicamycin treated cells using an anti-collagen-I antibody. Over 48 hours, both thapsigargin and tunicamycin abolished secretion of collagen-I (Fig. 2A). To examine if blockade of the secretory pathway had a similar effect on the secretion of collagen, cells were treated with brefeldin A and monensin (Fig. 2B and C). These treatments suppressed collagen secretion and induced ER stress, as shown by induction of CHOP transcription after 5 hours treatment (Fig. 2D). These effects were comparable to those of thapsigargin and tunicamycin, and likewise, both brefeldin A and monensin suppressed circadian fluctuations in Per2::luc cells (Fig. 2E).

**Figure 2:**
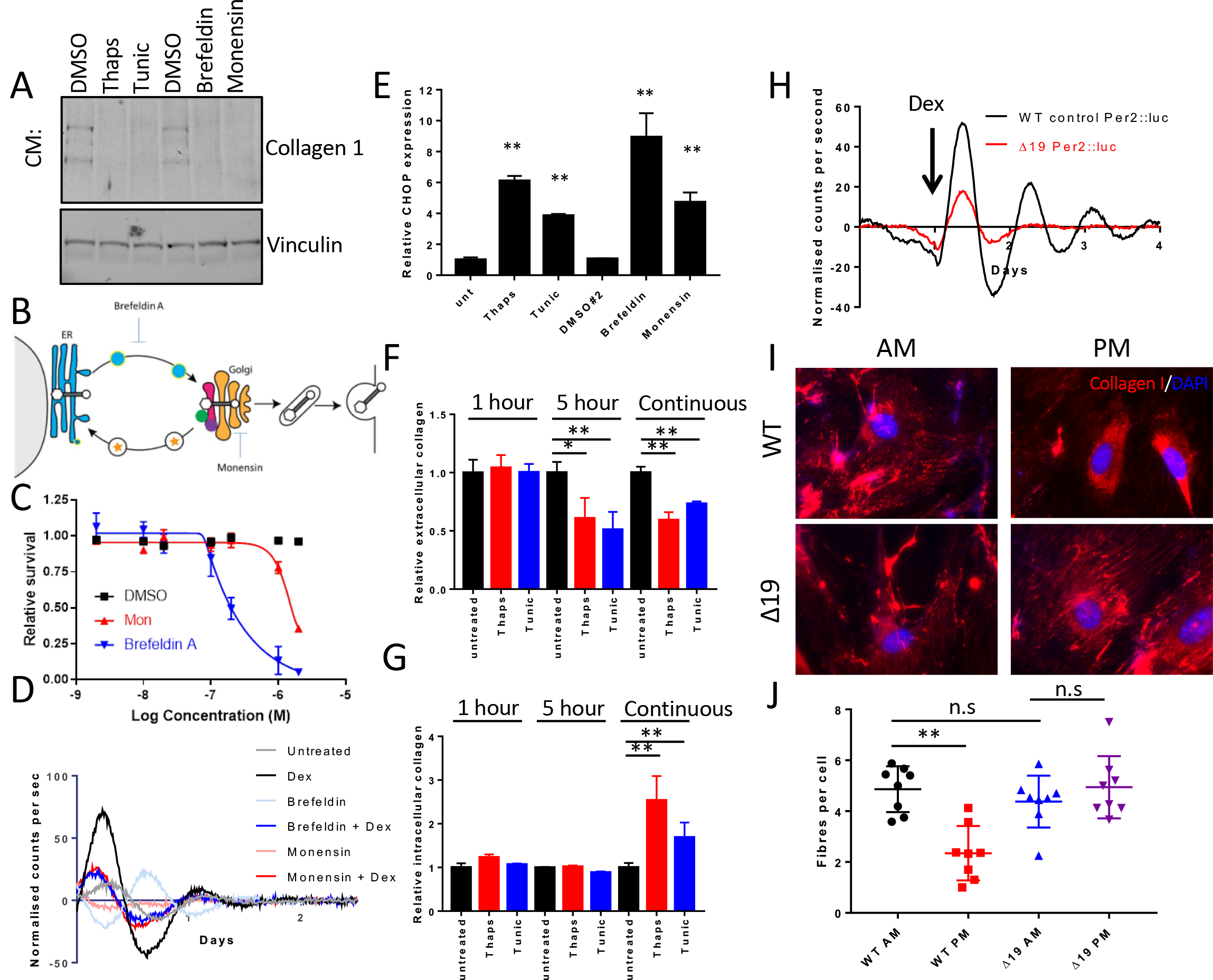
Impact of ER stress on collagen production. **A)** Western blot analysis on conditioned media (CM) collected from iTTFs after 48 hours of treatment with thapsigargin, tunicamycin, brefeldin, monensin suggests secreted collagen I is suppressed in treated cells. **B)** Diagram of the secretory pathway marking the position of action for the inhibitors brefeldin A and monensin. **C**) The effects of brefeldin A and monensin, at various concentrations, on the survival of fibroblasts following 72 hours treatment. **D**) The impact of 5 hours treatment of 20 nM brefeldin A and 10 nM monensin on Per2::luc signals in iTTFs. **E)** Levels of CHOP/DDIT3 indicate that the unfolded protein response has been activated following 5 hours treatment with brefeldin A and monensin, similar to the effects of thapsigargin and tunicamycin. The effects of thapsigargin and tunicamycin on the production of **F)** extracellular collagen and **G)** intracellular collagen, the effects of various treatment times are shown. **H**) Per2::luc signals in lung fibroblasts isolated from WT control and Clock Δ19 mice. Cells were recorded for 24 hours before treatment with 100 nM dexamethsone as Indicated. **I**) Collagen fibres formed by WT and ClockΔ19 fibroblasts 43 hours (AM) and 55 hours (PM) after synchronisation with dexamethasone. **J**) Scores of collagen fibres per cell from cultures in H. J) Transcript levels of ER stress related factors in WT and Clock Δ19 tendons * p=0.05 **p<0.01. paired t-test.

Having demonstrated that short term treatment with thapsigargin or tunicamycin can suppress circadian rhythm we sort to utilise this to assess the importance of circadian rhythm on the secretion and assembly of collagen fibres. Firstly, we showed that after treatment with 100 nM dexamethasone there is enhanced fibril assembly in iTTFs (Supplementary Fig. 4). We used a modified in-cell western approach to demonstrate the effects of thapsigargin and tunicamycin on collagen secretion. As expected, continuous treatment of fibroblasts with ER stress inducers reduced the assembly of extracellular collagen (Fig. 2F), which was accompanied by accumulation of intracellular collagen (Fig. 2G), in-line with reduced secretion of collagen into the medium (Fig. 2A). Having validated the in-cell western we then suppressed circadian rhythm using a 5-hour pulse of thapsigargin or tunicamycin prior to dexamethasone treatment. The results showed that disruption of rhythm suppressed the accumulation of extracellular collagen, after 72 hours, implicating circadian rhythms in coordinating collagen secretion and assembly. Importantly, the intracellular levels of collagen were unaffected with this treatment regime indicating that collagen had not been held in the ER. A one-hour treatment did not affect either extracellular or intracellular collagen. We have evaluated collagen fibre assembly in fibroblasts isolated from the ClockΔ19 mouse, which have a diminished circadian activity ^18^. These fibroblasts lack a sustained Per2::luc rhythm in response to dexamethasone treatment (Fig. 2H). In cultures of ClockΔ19 fibroblasts there were increased numbers of collagen fibres compared with wild type fibroblasts (Fig. 2I-J) and reduced ER stress related transcripts (Supplementary Fig.5), which supports recent observations that show increased collagen deposition in the tendons of ClockΔ19 mice ^4^. The deposition of collagen fibres by fibroblasts has also been found to be rhythmic ^4^. However, this rhythmicity is diminished in ClockΔ19 fibroblasts (Fig. 2I), which shows that circadian rhythm is required for normal collagen fibre homeostasis. Together these results suggest that ER stress can regulate circadian rhythm, and circadian rhythm affects collagen homeostasis, but collagen secretion *per se* does not require an intact circadian rhythm.

### BiP retains collagen in the ER but maintains circadian rhythm

Given the circadian rhythmicity of BiP protein levels, which peaks ahead of collagen-I (see Fig. 1A), we examined how BiP overexpression affects collagen-I secretion and the circadian rhythm. BiP was overexpressed in immortalised tail tendon fibroblasts (iTTFs) using retroviral transduction (Fig. 3A), which was confirmed by real-time PCR and western blotting (Fig. 3B and C). BiP overexpression led to inhibition of collagen fibre assembly at two time points following synchronisation (Fig. 3D and E). Both dexamethasone and serum shock induced a significant increase in collagen fibres in synchronised cultures (Supplementary Fig. 4) implying that synchronisation can aid the assembly of collagen fibres. Forskolin had the opposite effect (Supplementary Fig. 4); although forskolin can synchronize the circadian clock it also increases cAMP levels, which has previously been demonstrated to inhibit collagen trafficking ^19^ and TGFβ induced collagen production^20^. In control cells, fibre assembly was observed at times after BiP levels had peaked (see Fig. 1B). Overexpression of BiP did not alter the transcription of either *Col1a1* or *Col1a2* (Fig. 3F), suggesting that the reduced assembly of collagen fibres is a result of altered collagen transport through the secretory pathway. Immunofluorescence detection of intracellular collagen in BiP overexpressing (BiPoe) cells showed that the majority of the collagen co-localises with the ER marker protein disulphide isomerase (PDI), whereas in control cells, regions of the ER are clear from collagen. Thus, collagen is retained in the ER when BiP is overexpressed (Fig. 3G). The use of collagen hybridising peptide21 indicated that the collagen held in the ER of BiP over-expressing cells is largely unfolded (Supplementary Fig. 6). Therefore, we examined the impact of BiP over expression on circadian rhythm. To our surprise, the cells have a stronger inherent rhythm (Fig. 4A) and increased amplitude of Per2::luc signals following dexamethasone treatment (Fig. 4B and C). BiP is well known to suppress the activation of all three arms of the UPR; therefore, this result implies that the activation of the UPR arms is responsible for the suppression of circadian rhythms in cells undergoing ER stress. Real-time PCR analysis showed that both ATF6 and IRE1-XBP1 arms of the UPR are suppressed in BiP-overexpressing cells; however, there is enhanced activation of the ATF4 arm as shown by elevated CHOP expression (Fig. 4D).

**Figure 3:**
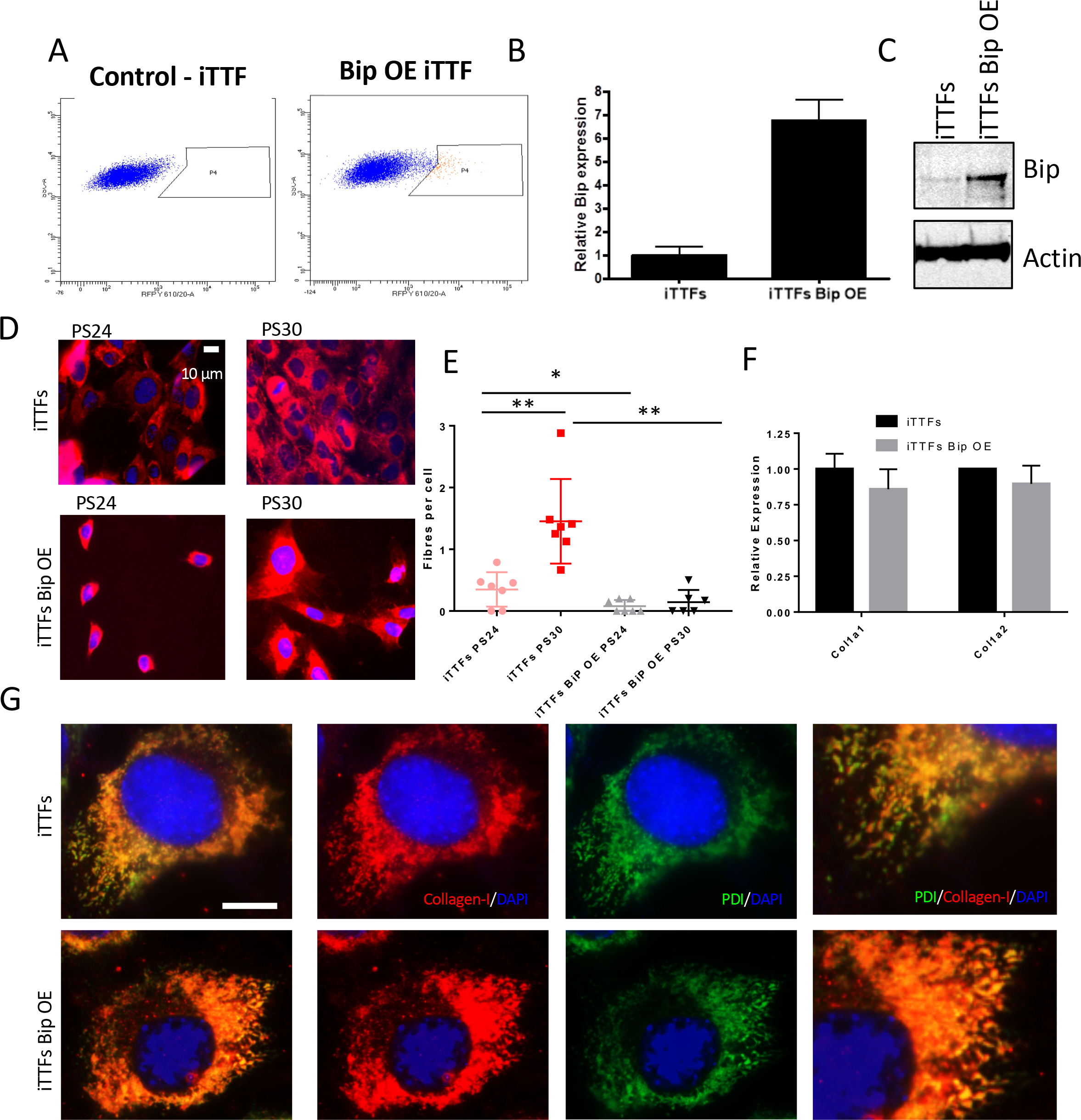
BiP retains collagen in the ER. **A)** Immortalised tail tendon fibroblasts were transduced with pCMMP-BiP-IRES-mRFP, RFP positive cells were sorted to form a BiP overexpressing cell line (iTTF + BiP). Levels of **B)** BiP mRNA and **C)** protein were assessed in sorted populations. **D)** Immunofluorescence detection of collagen fibres in control and BiP overexpressing immortalised tail tendon fibroblasts, cells were synchronised with dexamethasone and collected at different time points. PS24 and PS30, 24 hours and 30 hours post-synchronisation of the circadian clock by dexamethasone. **E)** the numbers of collagen fibres counted per cell at the indicated times. **F)** The effects of BiP overexpression on the transcript levels of type I collagen. **G)** Co-localisation of collagen I and the ER marker PDI in control and BiP overexpressing iTTFs. * p=0.05 **p<0.01 paired t-test.

**Figure 4:**
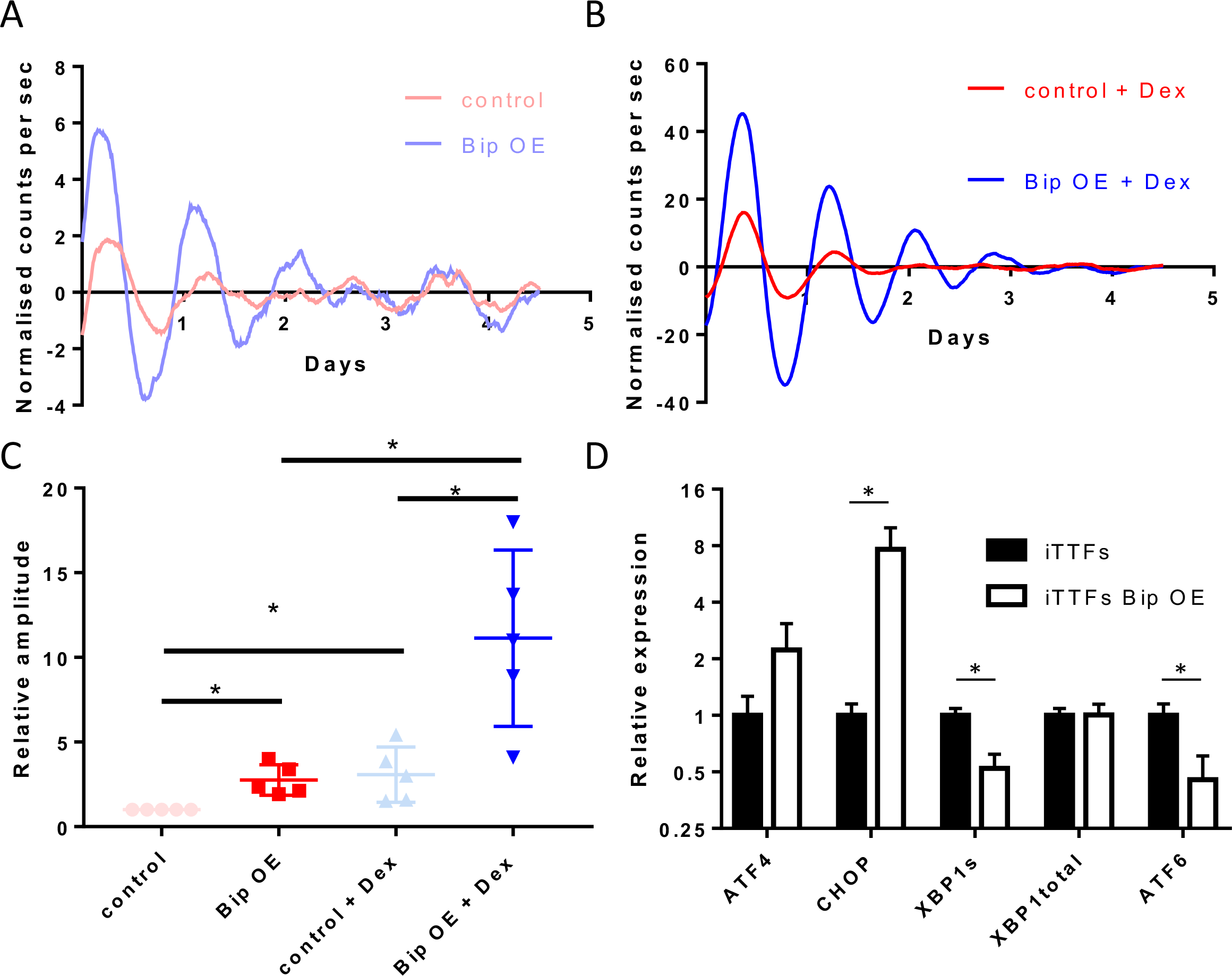
BiP overexpression strengthens circadian rhythm. **A)** Cells overexpressing BiP have a more robust circadian rhythm in unsynchronised populations and **B)** following dexamethasone-induced synchronisation. n=3, Representative traces are shown. **C)** Quantification of the amplitude of Per2::luc signals from A) and B) in the first 48 hours are quantified. **D)** Activation of the UPR in BiP overexpressing cells was assessed by monitoring the expression of ATF4/CHOP, Xbp1 splicing and ATF6. * p<0.05 paired t-test.

### Protein folding in the ER provides a checkpoint where circadian rhythm can be rapidly controlled

The influence of unfolded proteins on circadian clock regulation was explored using clinically-approved chemical chaperones, which aid the folding of proteins within the ER^22^. 4-phenylbutyric acid (4PBA) and ursodeoxycholic acid (UDCA) induced a visible rhythm in fibroblasts, but also greatly increased the amplitude of Per2::luc signals following dexamethasone treatment (Fig. 5A-F). These results resemble the effects observed with BiP overexpression and suggest that misfolded proteins are resident in the ER of cultured cells and through promoting their folding can rescue dampened circadian rhythms. The chaperone inhibitor 2-phenylethynesulfonamide (PES) had the opposite effect in suppressing the circadian rhythm, and was most likely due to the observed induction of the UPR (Supplementary Fig. 7). Assessment of the ER stress pathways following 4PBA treatment showed that there is suppression of both ATF6 and IRE1-XBP1 arms of the UPR, again mirroring the effects of BiP overexpression, and although ATF4 expression was enhanced, there was no elevated expression of CHOP (Fig. 5G). Treatment of tendon fibroblasts with 4PBA also led to enhanced secretion of collagen fibres (Fig. 5H and I) but without altering transcription of collagen-I (Fig. 5J). Taken together, these findings imply that protein folding in the ER is a rate limiting step in collagen biosynthesis and provides a checkpoint for control of the circadian clock.

**Figure 5:**
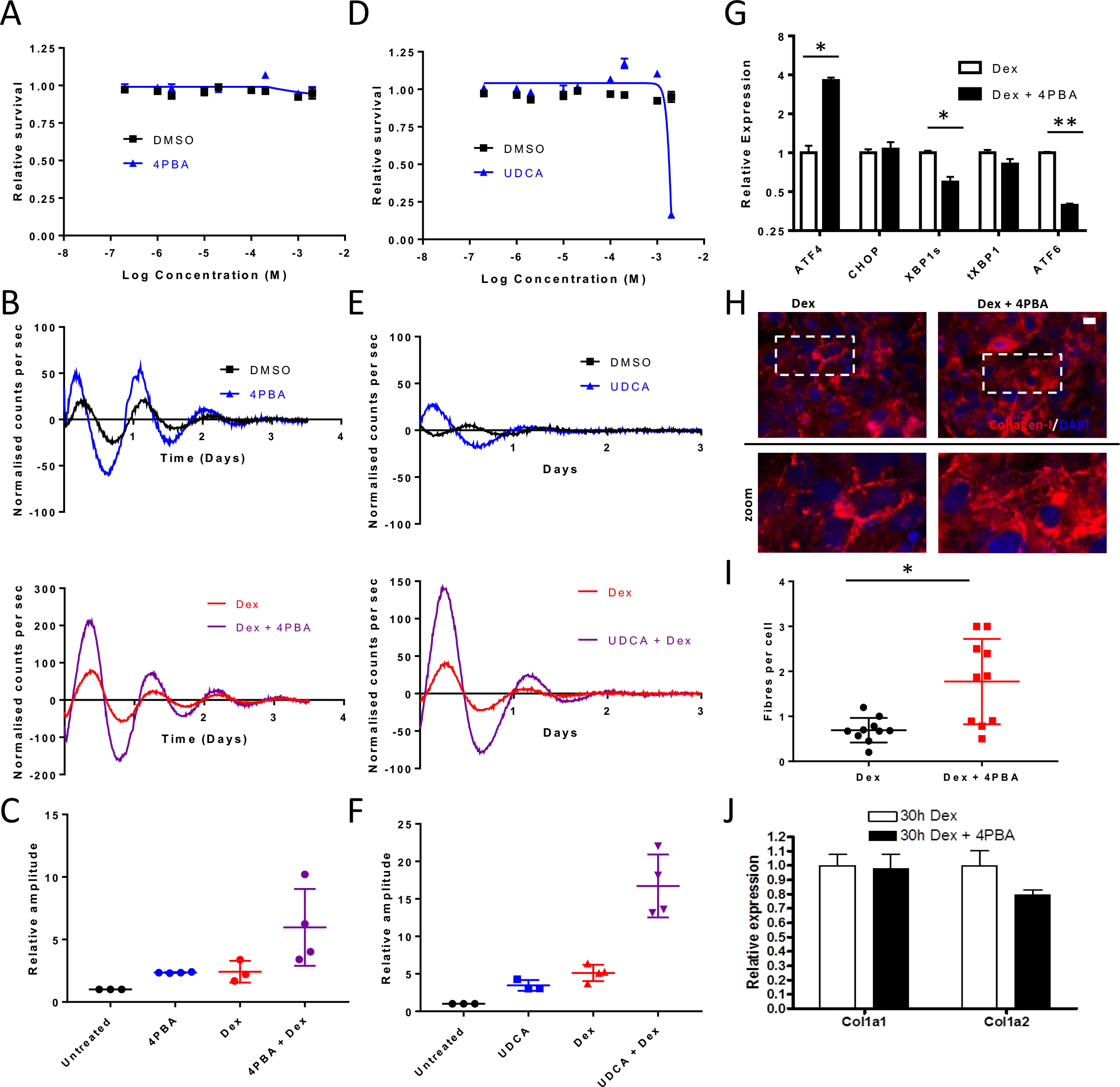
Chemical chaperones strengthen circadian rhythm and enhance collagen secretion. **A)** Effects of 4PBA treatment on the survival of immortalised tail tendon fibroblasts. **B)** The effects of 100 μM 4PBA on circadian rhythm in unsynchonised populations (upper panel) and following dexamethasone treatment (lower panel). **C)** Amplitudes of Per2::luc signals following 4PBA treatment. **D)** Effects of UDCA treatment on the survival of immortalised tail tendon fibroblasts. **E)** The effects of 100 μM UDCA on circadian rhythm in unsynchonised populations (upper panel), and following dexamethasone (lower panel). **F)** Amplitudes of Per2::luc signals following UDCA treatment. **G)** Effects of 4PBA on the expression of components of the unfolded protein response after 72 hours treatment. **H)** the effects of 4PBA treatment on the production of collagen fibres following 72 hours treatment, quantified in **I)**. **J)** The impact of 4PBA treatment on collagen I transcripts. * p<0.05, ** p<0.01 paired t-test.

## Discussion

The intrinsic circadian timing mechanism allows a cell to anticipate and adapt to daily rhythmic changes in physiological demand. In this study we have shown that BiP is rhythmic in tendon and cultured fibroblasts, and that the peak of BiP protein expression occurs just ahead of the peak of procollagen-I expression. We further showed that ER stress, caused by inhibiting SERCA (using thapsigargin) or expression of misfolded proteins in the ER, eliminates the circadian rhythm and decreases collagen secretion. Whilst thapsigargin is an irreversible inhibitor of SERCA, we found that by removing thapsigargin from cells, circadian rhythms could be reinitiated. This suggests that the loss of response to clock synchronising cues is due to the presence of misfolded protein. A recent study by Walton *et al* demonstrated a similar degree of circadian rhythm suppression under hypoxic conditions; whilst not directly addressed in their report, a strong induction of genes involved in the UPR was observed^23^. Hypoxia induction is well known to induce protein misfolding and activation of the UPR^24, 25^, again implicating ER stress control of circadian rhythm.

The fluctuations in BiP levels appear to play a protective role in maintaining circadian rhythms, as BiP overexpression led to a more robust circadian rhythm, presumably because of the saturation of all three sensors of the UPR. Thus, we are proposing that the circadian clock has a role in pre-emptively preventing ER stress in normal cells, by upregulating BiP in anticipation of a surge in protein translation. However, similar to what has been observed for other proteins ^26, 27^, overexpression of BiP led to retention of procollagen in the ER, and the procollagen was in a non-helical conformation concomitant with reduced assembly of collagen fibres. A balance is therefore struck to have sufficient BiP to allow saturation of the UPR sensors and a robust circadian rhythm, but not too much to inhibit folding of procollagen.

Yeung *et al.* showed that tendon tissue contains has an autonomous circadian clock. In their time-series microarray study they identified 745 rhythmic transcripts that had a 24-hour oscillation. However, neither *Col1a1* nor *Col1a2*, or indeed any transcripts encoding collagens, were rhythmic^28^. In a subsequent study, Yeung and Garva *et al.* demonstrated that the levels of readily-extractable collagen-I oscillate with a 24 hour period, with the majority of procollagen synthesis occurring in the evening in mouse tail and Achilles tendon^4^. The circadian rhythmicity of collagen-I is therefore generated in the ER, rather than through transcriptional regulation. The ramp in procollagen concentration in the ER at night might be expected to lead to ER stress if the levels of nascent unfolded protein are allowed to escalate unchecked. However, levels of BiP are also circadian-regulated and peak a few hours ahead of the peak of procollagen. This suggests that the circadian clock prepares the cell for the surge in nascent procollagen chains being synthesised by ramping the synthesis of BiP. After the surge of procollagen has passed, BiP levels reduce to avoid swamping the ER stress sensors and thereby re-establish homeostasis to the UPR sensing machinery. We have demonstrated here that induction of ER stress can ablate the circadian rhythm in normal fibroblasts, suggesting that this surge of BiP may influence how cells perceive external circadian cues. The fluctuations in BiP levels appear to play a protective role by maintaining circadian rhythms in normal cells.

BiP overexpression, and chemical chaperones treatment, led to suppression of Xbp1 splicing and ATF6 transcription, and at the same time enhancing circadian rhythm, implying that these arms of the UPR are integrated into the feedback control of the molecular clock during ER stress. The ATF4 arm was largely unaffected. It has previously been demonstrated that ATF4 knockout MEFs have suppressed amplitude of Per2::luc rhythms ^29^, suggesting this pathway is essential for the robust rhythms, and may further explain why these pathways remain active whilst the other arms of the UPR are suppressed. Per1 mRNA was shown to be a target for the exonuclease activity of IRE1, and its activation is associated with reduced Per1 expression^30^, presenting a potential feed-in point for controlling the circadian circuitry. In a genome-wide siRNA screen to identify modulators of circadian rhythm, ATF6 siRNA was identified to shorten the circadian period in U2OS cells^31^ and is also another potential input point. Here, we did not observe a shortening of period as a result of ER stress induced ATF6 expression; however this is in-line with a recent study which demonstrated a 10 hour shift in circadian rhythm in cancer cells undergoing ER stress^14^. We have shown that ER stress or misfolded proteins can dramatically suppress clock activation in normal fibroblasts, and this poses the question that cancer cells may negate this suppressive role of the UPR on the clock in order to survive. In normal cells undergoing ER stress, a decision of whether the stress is resolved or the cell dies has to be made. It is plausible that by uncoupling the cell from circadian control, this decision can be made more efficiently. It has been proposed that cancer cell survival is enhanced through suppression of Clock and Bmal1 ^14^, however, this may result in survival even when misfolded proteins have not been eliminated. Both ER stress and UPR activation are well documented in many cancers implicating accumulation of misfolded proteins ^32^. Similarly, ER stress is active in patients with idiopathic pulmonary fibrosis ^33^, liver fibrosis ^34^ and kidney fibrosis ^35, 36^. Given that activation of ER stress would normally lead to suppression of circadian rhythms and collagen synthesis, this suggests that cells within diseased tissues have circumvented this physiological control in order to drive fibrosis. Establishing how this is achieved in diseased tissues is the focus of future studies.

## Methods

### Cell isolation and culture

Tail tendon fibroblasts were released from 6-8 week PER2::Luc C57BL/6J mouse (kindly donated by Professor Joseph Takahashi of the University of Texas Southwestern Medical Center) tail tendons by 4 mg/ml bacterial collagenase type 4 (Worthington Biochemical Corporation) in EDTA-free trypsin (2.5 g/l) as previously described ^37^. Cells were cultured in complete media (Dulbecco’s modified Eagle’s medium: Nutrient Mixture F-12 containing 4500 mg/L glucose, L-glutamine, and sodium bicarbonate, supplemented with 10% foetal calf serum) at 37 °C, in 5% CO_2_. Immortalized lines were generated by retroviral expression of mTERT ^38^ (iTTF), similarly to generate BiP overexpressing cells were we utilised pCMMP-BiP-IRES-mRFP a gift from Bill Sugden (Addgene plasmid #36975)^39^, using methods previously described^40^. BiP overexpressing cells were flow sorted based on mRFP expression. Stable cell lines expressing mutant thyroglobulin were generated by transfecting pcDNA3-neo-ColII-TgCOG into iTTFs using Fugene 6 (Promega). Forty-eight hours after transfection cells were maintained in 200 μg/mL neomycin for 2 weeks, prior to analysis of luminescence neomycin was removed from culture medium, as controls cells transfected with pEGFP-N1 (Clontech) and selected as above.

### Luminometry

LumiCycle apparatus (Actimetrics) was used for real-time quantitative bioluminescence recording. Tail tendons, 10 mg, were placed in 30 mm cell culture inserts (0.4 μm pore size, Millipore) inside 35 mm dishes in recording medium (DMEM without phenol-red (Sigma, D2902), supplemented with 4 g/L glucose, 5% FCS, HEPES, sodium bicarbonate and 0.1 mM Luciferin). iTTFs were seeded 24 hours, as described above, prior to culture in recording medium. 100 nM dexamethasone was added for 30 minutes to synchronise the circadian rhythms. Baseline subtraction was carried out using a 24-hr moving average. Amplitude was calculated as peak-trough difference in bioluminescence of the first and second peak, as indicated, using base-line subtracted data.

### Western Blot

Proteins were extracted using urea buffer and analysed by western blotting as previously described ^41^. Primary antibodies used were mouse mAb to BiP (1:1000; sc-376768, Santa Cruz), rabbit pAb to Collagen-I (1:500; OARA02579, Gentaur), mouse mAb to GAPDH (1:10000; clone GAPDH-71.1, Sigma), mouse mAb to vinculin (1:800; V9131, Sigma), mouse mAb to beta-actin (1:2000; sc-8432, Santa Cruz).

### Dose response curves

The appropriate dosages of thapsigargin and tunicamycin for *in vitro* treatments were determined in triplicate using the AlamarBlue assay in 96 well format ^42^.

### Real time PCR

For qPCR analysis primers designed using Assay Design center (Roche, Germany), the sequences used are indicated in Table 1. Reactions were run using Power-UP Sybr Master mix (Applied Biosystems) with cDNA generated using Taqman reverse transcription kit (Applied Biosystems). 0.5 μg RNA extracted using Trizol (Life Technologies) was used to generate cDNA which was then diluted 20-fold for PCR. PCR reaction were run on the CFX96 Real time system (BioRad). Target transcript expression was normalised to the geometric mean of Rplp0, Gapdh and Actb.

### Immunofluorescence and in-cell western

For immunofluorescence analysis ells were fixed with 4% paraformaldehyde and permeabilized with 0.5% Triton-X/PBS. Collagen-I was detected using a rabbit pAb (1:200; OARA02579, Gentaur), and ER was identified using a mouse mAb against protein disulphide isomerase (Pdi) (1:100; ab190883, Abcam). Images were collected on a Leica TCS SP5 AOBS inverted confocal using a 60× / 0.50 Plan Fluotar objective and 3× confocal zoom. The confocal settings were as follows, pinhole 1 airy unit, scan speed 1000 Hz unidirectional, format 1024 × 1024. Images were collected using PMT detectors with the following detection mirror settings; FITC 494-530 nm; Cy5 640-690 nm using the 488 nm (20%), 594 nm (100%) and 633 nm (100%) laser lines respectively. To eliminate crosstalk between channels, the images were collected sequentially.

When acquiring 3D optical stacks the confocal software was used to determine the optimal number of Z sections. Only the maximum intensity projections of these 3D stacks are shown in the results. For imaging non-helical collagen, cells were permeabilized as above and stained with anti-Collagen-I (1:200; OARA02579, Gentaur) and 20 μM 5-FAM conjugated collagen hybridizing peptide (F-CHP, 3Helix). Cells were stained for 2 hours at 4 °C. Images were collected on a Zeiss Examiner A1 upright microscope using a 63× / 1.4 N-Acroplan objective and captured using a Coolsnap ES camera (Photometrics) through Metavue v7.8.0.0 software (Molecular Devices). Specific band pass filter sets for DAPI, FITC and Cy3 were used to prevent bleed through from one channel to the next. For in-cell westerns, iTTF were grown in 96 well plates in the presence of 200 μM ascorbic acid, 24 hours after plating cells were treated with tunicamycin and thapsigargin as indicated before removing treatment and adding full medium supplemented with ascorbic acid and dexamethasone. After 72 hours cells were fixed with 4% PFA. To assess extracellular collagen cells were directly stained with an anti-collagen antibody (1:1000; OARA02579, Gentaur) overnight, which as detected using donkey anti-rabbit 800 antibody (1:10,000, #5151s, Cell signal technologies), each well was counterstained with the cell permeating dye DRAQ5. The plate was scanned using the Odyssey CLx (Li-Cor). The collagen signal quantified following normalisation to DRAQ5. To assess intracellular collagen the plates were permeabilised as for immunofluorescence studies before being stained and quantified in the same manner as extracellular collagen.

## Acknowledgements

We would like to thank Raymond Boot-Handford for expression vectors. The research was funded by Wellcome Trust Investigator and Wellcome Centre Core Awards to K.E.K. (110126/Z/15/Z and 203128/Z/16/Z). P.A. is funded by NIH DK40344. B.C. is funded by a Wellcome 4-year PhD studentship (210062/Z/17/Z). Light microscopes in the Bioimaging Facility were additionally supported by the University of Manchester Strategic Fund.

